# Chondrocytes in the resting zone of the growth plate are maintained in a Wnt-inhibitory environment

**DOI:** 10.1101/2020.11.15.383331

**Authors:** Shawn A. Hallett, Wanida Ono, Yuki Matsushita, Naoko Sakagami, Koji Mizuhashi, Nicha Tokavanich, Mizuki Nagata, Annabelle Zhou, Takao Hirai, Henry M. Kronenberg, Noriaki Ono

## Abstract

Chondrocytes in the resting zone of the postnatal growth plate are characterized by slow cell cycle progression, and encompass a population of parathyroid hormone-related protein (PTHrP)-expressing skeletal stem cells that contribute to the formation of columnar chondrocytes. However, how these chondrocytes are maintained in the resting zone remains undefined. We undertook a genetic pulse-chase approach to isolate slow cycling, label-retaining chondrocytes (LRCs) from the growth plate using a chondrocyte-specific doxycycline-controllable Tet-Off system regulating expression of histone 2B-linked GFP. Comparative RNA-seq analysis identified significant enrichment of inhibitors and activators for Wnt/β-catenin signaling in LRCs and non-LRCs, respectively. Activation of Wnt/β-catenin signaling in PTHrP^+^ resting chondrocytes using *Pthrp-creER* and *Apc*-floxed allele impaired their ability to form columnar chondrocytes. Therefore, slow-cycling chondrocytes are maintained in a canonical Wnt-inhibitory environment within the resting zone, unraveling a novel mechanism regulating maintenance and differentiation of PTHrP^+^ skeletal stem cells of the postnatal growth plate.

## Introduction

The epiphyseal growth plate, a disk of cartilaginous tissues with characteristic columns of chondrocytes formed between the primary and secondary ossification centers, is an innovation of amniotes (reptiles, birds and mammals) that facilitates explosive endochondral bone growth (Wuelling & Vortkamp, 2019). The postnatal growth plate is composed of three morphologically distinct layers of resting, proliferating and hypertrophic zones, in which chondrocytes continue to proliferate well into adulthood, especially in mice, therefore functioning as the engine for endochondral bone growth (Hallett et al., 2019; Kronenberg, 2003). By adulthood, a large majority of hypertrophic chondrocytes undergo apoptosis or transdifferentiate into osteoblasts, marking the completion of the longitudinal growth phase and skeletal maturation (Roach et al., 1995; Yang et al., 2014; Zhou et al., 2014).

Of the three layers, the resting zone has two important functions in maintaining the growth plate. First, early studies postulated that resting chondrocytes feed their daughter cells into the adjacent proliferating zone and contribute to longitudinal growth of postnatal endochondral bones (Hunziker, 1994). More recently, the resting zone has been established as a niche for skeletal stem cells, initially demonstrated by surgical auto-transplantation experiments in rabbits (Abad et al., 2002), and subsequently by lineage-tracing experiments in mice (Mizuhashi et al., 2018; Newton et al., 2019). Second, chondrocytes in the resting zone express parathyroid hormone-related protein (PTHrP) that maintains proliferation of chondrocytes in a cell non-autonomous manner and delays their hypertrophic differentiation, thus sustaining longitudinal growth (E. et al., 1997). The proliferating zone is concertedly maintained by PTHrP released from the resting zone and Indian hedgehog (Ihh) synthesized by pre-hypertrophic chondrocytes; the proliferating zone in turn provides instructive cues to regulate cell fates of PTHrP^+^ chondrocytes (Mizuhashi et al., 2018). Therefore, the resting zone functions as a critical constituent of the tight feedback system (the PTHrP–Ihh feedback loop) that maintains growth plate structures and longitudinal bone growth. However, mechanisms regulating self-renewal and differentiation capabilities of resting zone chondrocytes remain largely unknown.

In this study, we set out to undertake an unbiased approach to define the molecular mechanism regulating maintenance and differentiation of chondrocytes in the resting zone (‘slow-cycling chondrocytes’). To achieve this goal, we developed a chondrocyte-specific genetic label-retention strategy to isolate slow-cycling chondrocytes from the postnatal growth plate. Our comparative transcriptomic analysis revealed unique molecular signatures defining the characteristics of slow-cycling chondrocytes, with particular enrichment for inhibitors of the canonical Wnt signaling pathway. Subsequent functional validation based on a cell-lineage analysis identified that, when Wnt/β-catenin signaling was activated, PTHrP^+^ resting chondrocytes were decreased in number during initial formation and established columnar chondrocytes less effectively in the subsequent stages. These data lead to a new concept that PTHrP^+^ skeletal stem cells may be maintained in a canonical Wnt inhibitory environment within the resting zone niche of the postnatal growth plate.

## Results

### 1.1. A genetic label-retention strategy to identify slow-cycling chondrocytes

Chondrocytes in the resting zone of the postnatal growth plate (‘resting’ or ‘reserve’ chondrocytes) are characterized by their slow cell cycle progression that is much slower than that of chondrocytes in the proliferating zone. As a result, these slow-cycling chondrocytes retain nuclear labels much longer than their more rapidly dividing progeny in the proliferating zone, which are therefore termed as label-retaining chondrocytes (LRCs) (Walker & Kember, 1972). First, we undertook a genetic approach to fluorescently isolate LRCs from the growth plate based on a chondrocyte-specific pulse-chase protocol. To this end, we first generated transgenic mice expressing a tetracycline-controlled transactivator under a *Col2a1* promoter (hereafter, Col2a1-tTA), and combined this line with a *Col1a1* locus harboring a Tet-responsive element (TRE)-histone 2B-bound EGFP (H2B-EGFP) cassette (hereafter, TRE-H2B-EGFP) (Fig. 1a, left panel). In this Tet-Off system, tTA binds to TRE in the absence of doxycycline and activates H2B-EGFP transcription (pulse), whereas tTA dissociates from TRE in the presence of doxycycline, shutting off H2B-EGFP transcription (chase) (Fig. S1a). In Col2a1-tTA; TRE-H2B-EGFP mice, *Col2a1*^*+*^ chondrocytes accumulate H2B-EGFP in the nucleus without doxycycline, and upon initiation of doxycycline feeding, *de novo* transcription of *H2B-EGFP* mRNA becomes suppressed. After a long chase period, H2B-EGFP is preferentially diluted in highly proliferating cells and their progeny, whereas slow-cycling cells retain a high level of the label, marking them as LRCs.

**Figure 1.**
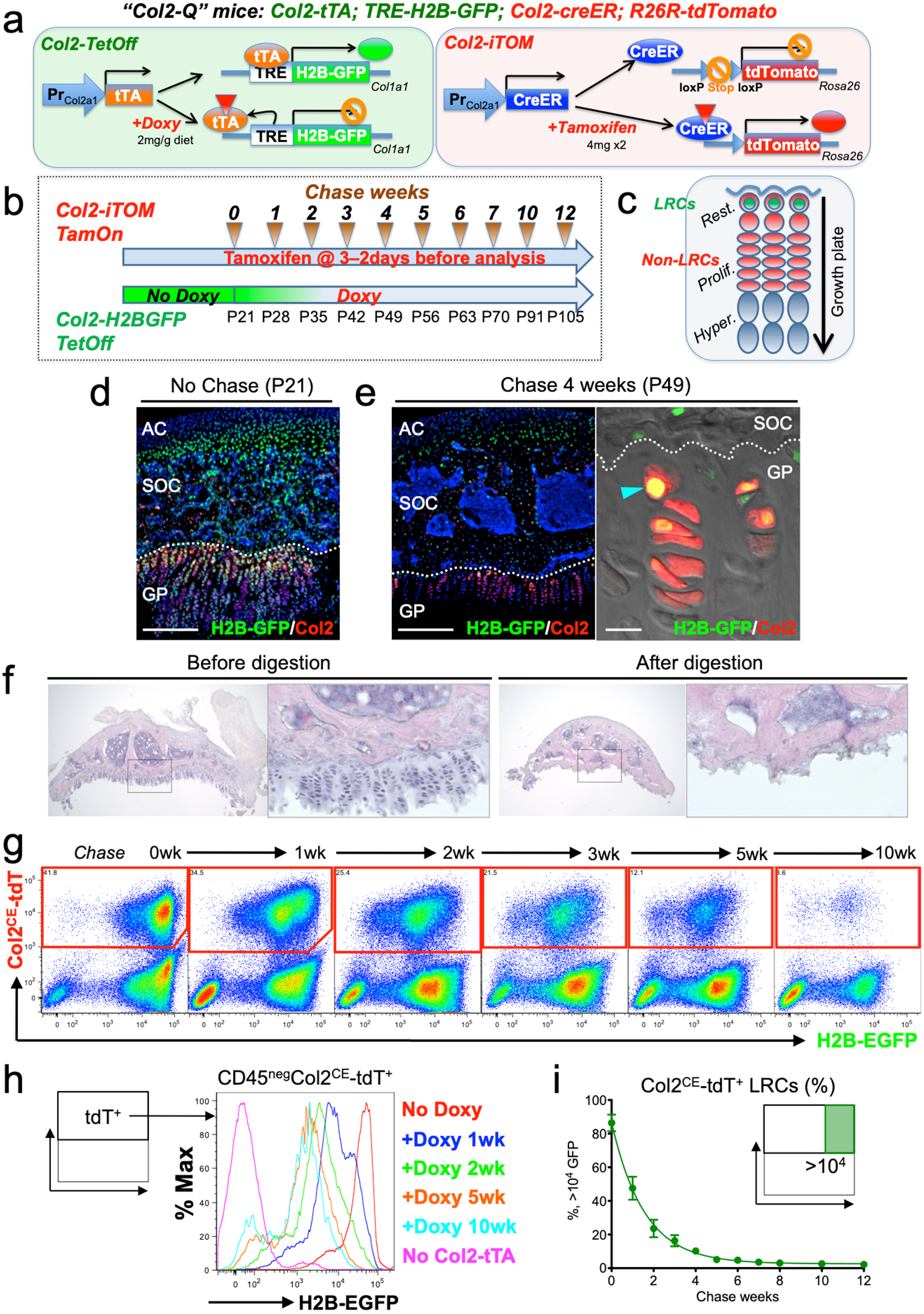
A double-color genetic label-retaining strategy to identify and isolate slow-cycling chondrocytes of the growth plate. **(a)** “Col2-Q” quadruple transgenic system composed of two chondrocyte-specific bigenic Col2-Tet-Off (*Col2a1-tTA*; *TRE-H2B-EGFP*) and Col2-iTOM (*Col2a1-creER*; *R26R-tdTomato*) systems. H2B-EGFP expression can be shut off by doxycycline diet (2mg/g diet), while tdTomato expression can be induced by two doses of tamoxifen (4mg) administered shortly prior to analysis (3 and 2 days before). **(b)** Experimental design to identify label-retaining chondrocytes (LRCs) in the growth plate. Col2-Q mice are fed with doxycycline (Doxy) starting from postnatal day (P) 21 (Chase). The mice are analyzed after the indicated number of weeks; at each time point, two doses of tamoxifen are administered shortly before analysis to induce tdTomato expression. **(c)** Diagram for predicted outcomes. LRCs are expected to retain green nuclei with red cytoplasm, and located at the resting zone. Rest.: resting zone, Prolif.: proliferating zone, Hyper.: hypertrophic zone. **(d,e)** Col2-Q distal femur growth plates with tamoxifen injection shortly before analysis. (d): No chase, without Doxy at P21. (e): After 4 weeks of chase, on Doxy for 4 weeks at P49, right panel: high-power confocal image. Arrowhead: label-retaining chondrocytes. AC: articular cartilage, SOC: secondary ossification center, GP: growth plate. Dotted line: border between growth plate and secondary ossification center. Blue: DAPI, grey: DIC. Scale bars: 500μm, 20μm (confocal in e). *n*=3 mice at each time point. **(f)** Epiphysis of proximal tibia, before and after serial collagenase digestions. Right panels: magnified views of the dotted areas showing growth plate. *n*=3 mice at each step. **(g-i)** Flow cytometry analysis of dissociated Col2-Q growth plate cells. (g): Pseudo-color plots of CD45^neg^ cells at the indicated number of weeks in chase. Red gates: Col2a1-creER/tdTomato^+^ (Col2^CE^-tdT^+^) cells. (h): Histogram of CD45^neg^Col2^CE^-tdT^+^ cells showing the distribution of H2B-EGFP^+^ cells as the percentage of the maximum count. Red line: P21 (No Doxy), blue line: P28 (+Doxy 1wk), green line: P35 (+Doxy 2wk), orange line: P56 (+Doxy 5wk), light blue line: P91 (+Doxy 10wk), pink line: No Col2-tTA control at P21. (i): Percentage of >10^4^ H2B-EGFP^+^ LRCs among total Col2^CE^-tdT^+^ cells. *x* axis: weeks in chase, *y* axis: % of cells > 10^4^ unit of GFP. *n*=9 mice (0 week, 1 week), *n*=7 mice (2 weeks, 5 weeks), *n*=6 mice (3 weeks, 4 weeks), *n*=5 mice (6 weeks) and *n*=3 mice (7 weeks, 8 weeks, 10 weeks, 12 weeks). Data are presented as mean ± s.d.

In order to evaluate the labeling efficiency of the system, we first analyzed double heterozygous Col2a1-tTA/+; TRE-H2B-EGFP/+ mice at postnatal day (P) 7 and P21 without doxycycline. While most of chondrocytes in the growth plate were marked by a high level of H2B-EGFP at P7 (Fig. S1b), fewer than half of columnar chondrocytes in the growth plate were marked by H2B-EGFP at P21 (Fig. S1c), demonstrating the inefficiency of the Tet system in postnatal growth plate chondrocytes. To circumvent this problem, we further generated double homozygous Col2a1-tTA/Col2a1-tTA; TRE-H2B-EGFP/TRE-H2B-EGFP mice and analyzed these mice at P21 without doxycycline. A great fraction of columnar chondrocytes was marked by H2B-EGFP (Fig. S1d), indicating that the labeling efficiency can be improved in a transgene dosage-dependent manner in this system.

Subsequently, we tested the effectiveness of this chondrocyte-specific Tet-Off system by pulse-chase experiments. We fed double heterozygous Col2a1-tTA/+; TRE-H2B-EGFP/+ mice with doxycycline for 5 weeks starting from P21 to shut off de novo H2B-EGFP expression. We started the chase at P21 because the secondary ossification center was fully developed within the epiphysis by this time. After the chase, the H2B-EGFP signal was largely abrogated in the growth plate region, with only a small fraction of cells in the resting zone near the top of the growth plate retaining H2B-EGFP (Fig. S1e, arrowheads) expression. However, we also noticed that a low level of H2B-EGFP signal persisted in adjacent osteoblasts and osteocytes in the epiphysis even after the chase (Fig. S1e, arrows), making it difficult to distinguish LRCs from these cells. Analysis of TRE-H2B-EGFP/+ mice without a Col2a1-tTA transgene at P28 revealed that osteoblasts and osteocytes expressed a low level of H2B-EGFP (Fig. S1f, arrows). These findings indicate that LRCs can be identified within the top of the growth plate by a chondrocyte-specific Tet-Off system regulating H2B-EGFP expression, although these cells cannot be easily distinguished from adjacent osteoblasts and osteocytes solely based on fluorescent intensity in histological sections.

### 1.2. Col2-Q system: A double-color quadruple transgenic strategy to identify LRCs in the growth plate

To circumvent the technical issues hampering isolation of LRCs from the growth plate resting zone, we further included a *Col2a1-creER* transgene that activates an *R26R-tdTomato* reporter in a tamoxifen-dependent manner, as a means to specifically mark growth plate chondrocytes (M. Chen et al., 2007). We generated quadruple homozygous transgenic mice – “Col2-Q” mice: *Col2a1-tTA*; *TRE-H2B-EGFP*; *Col2a1-creER*; *R26R-tdTomato* (Fig. 1a), and treated these mice with tamoxifen (4 mg) twice shortly before analysis (3 and 2 days before analysis, “short protocol”) to obtain Col2a1-creER-tdTomato^+^ cells (hereafter, Col2^CE^-tdT^+^ cells). After the pulse-chase protocol with doxycycline, LRCs are expected to be identified as cells with green nuclei and red cytoplasm, which are localized in the resting zone of the growth plate (Fig. 1b,c). First, we analyzed Col2-Q mice at P21 without doxycycline (“No Chase”). A great majority of cells in the epiphysis, including those in the growth plate and the secondary ossification center, but not on the articular surface, were H2B-EGFP^high^ (Fig. 1d, cells with green nuclei). This short protocol of tamoxifen injection marked a great number of chondrocytes in the growth plate, but not in the articular cartilage (Fig. 1d), indicating that this double-color strategy can effectively identify H2B-EGFP^high^ growth plate chondrocytes at this stage. Second, Col2-Q mice were fed with doxycycline from P21 to shut off new H2B-EGFP synthesis for 4 weeks (chase) and were then treated with the short protocol of tamoxifen injection. After the chase, LRCs were identified at a specific location near the top of the growth plate in the resting zone, retaining a higher level of H2B-EGFP signal (Fig. 1e). In addition, most of these H2B-EGFP^high^ cells in the growth plate were simultaneously marked as Col2^CE^-tdT^+^ (Fig. 1e, right panel, arrowhead), while more rapidly dividing and morphologically distinct columnar chondrocytes were not marked by H2B-EGFP signal. Therefore, our Col2-Q quadruple transgenic strategy can effectively mark LRCs primarily in the resting zone of the postnatal growth plate.

### 1.3. A flow cytometry-based identification and isolation of LRCs from Col2-Q mice

We next established a protocol to harvest chondrocytes from the postnatal growth plate. We manually removed epiphyses from four long bones (bilateral distal femurs and proximal tibias [Fig. 1f, left panel, shown is a dissected epiphysis from a tibia]). With this protocol, the growth plate was sheared at the hypertrophic layer with the remainder attached to the epiphysis. We further digested dissected epiphyses serially with collagenase to release these cells into single-cell suspension. Five rounds of digestion completely liberated cells from the growth plate, while cells on the articular surface were almost undigested (Fig. 1f, right panel). Subsequently, we used a flow cytometric approach to analyze single cells dissociated from the Col2-Q postnatal growth plate at sequential time points before and after the chase, particularly in a CD45-negative non-hematopoietic fraction. Col2a1^CE^-tdT^+^ cells were clearly distinguishable from unlabeled cells at all time points investigated (Fig. 1g). Before the chase started at P21 (therefore without doxycycline feeding), 86.4±5.0% of Col2a1^CE^-tdT^+^ cells retained >10^4^ units of H2B-EGFP (Fig. 1g, leftmost panel). The fraction of a label-retaining population (GFP^high^, retaining >10^4^ unit of H2B-EGFP signal) within a Col2^CE^-tdT^+^ population gradually decreased as the chase period extended (Fig. 1g). These plots fit into a non-linear decay curve (Y0: 86.5±1.3%; Plateau: 2.6±0.9%; T^1/2^=0.99~1.18 week) (Fig. 1i). Virtually no GFP^+^ cells were observed in the absence of a Col2a1-tTA transgene (Fig. 1h, magenta line), while levels of GFP^+^ cells were maintained from five to ten weeks of chase (Fig. 1h, orange and teal lines). Therefore, these findings demonstrate that LRCs can be effectively identified and isolated from postnatal Col2-Q growth plates by combined microdissection, enzymatic digestion and flow cytometry-based approaches.

## 2. A comparative RNA-seq analysis reveals a unique molecular signature of LRCs

Subsequently, we isolated slow-cycling chondrocytes using fluorescence-activated cell sorting (FACS) at a 4 week-chase time point, when the GFP^high^ label-retaining fraction (>10^4^ unit) was sufficiently enriched (Fig. 1i). In this experiment, LRCs were defined as GFP^high^ cells retaining H2B-EGFP signal at the top 10% brightness (*x* > 10^4^ unit), whereas non-LRCs were defined as other GFP^mid-low^ cells (10^3^ < *x* < 10^4^ unit) (Fig. 2a). Cells were collected from multiple littermate mice for each of three independent experiments. To assess RNA quality, we conducted an RNA Integrity Number (RIN) assay (Schroeder et al., 2006) from total RNAs isolated from LRCs and non-LRCs. Cellular RNA levels from each population had sufficient quality for downstream application (RIN>8.0, Fig. 2b), which were further subjected to amplification and deep sequencing. An unsupervised clustering analysis demonstrated that LRCs and non-LRCs biological triplicate samples each clustered together (Fig. 2c), indicating that slow-cycling chondrocytes in the postnatal growth plate possess a biologically unique pattern of transcriptomes compared to more rapidly diving non-LRCs. Analyses of differentially expressed genes (DEGs) revealed that 799 genes were differentially expressed between the two groups (fold change ≥ ± 2), of which 427 and 372 genes were upregulated in LRCs and non-LRCs, respectively (Fig. 2d). Representative genes upregulated in LRCs included known markers for resting chondrocytes, such as *Pthlh* (also known as *Pthrp*, x2.6)(X. Chen et al., 2006) and *Sfrp5* (x2.4); (Chau et al., 2014; Lui et al., 2010) in addition to novel markers, such as *Gas1* (x12), *Spon1* (x10) and *Wif1* (x3.8). Similarly, representative genes upregulated in non-LRCs included both known and novel markers for proliferating and pre-hypertrophic chondrocytes, such as *Ihh* (St-Jacques et al., 1999) (x54), *Col10a1* (Gu et al., 2014) (x11), *Mef2c* (Arnold et al., 2007) (x5.1), *Pth1r* (Hirai et al., 2011) (x3.0), *Sp7* (Nakashima et al., 2002) (x2.4) and *Dlx5* (Robledo et al., 2002) (x2.2) as well as *Clec11a* (Yue et al., 2016) (x2.9) and *Cd200* (Etich et al., 2015) (x2.1). Therefore, these identified enriched genes support the precision and accuracy of comparative RNA-seq analysis of LRCs and non-LRCs isolated by cell sorting from the growth plate.

**Figure 2.**
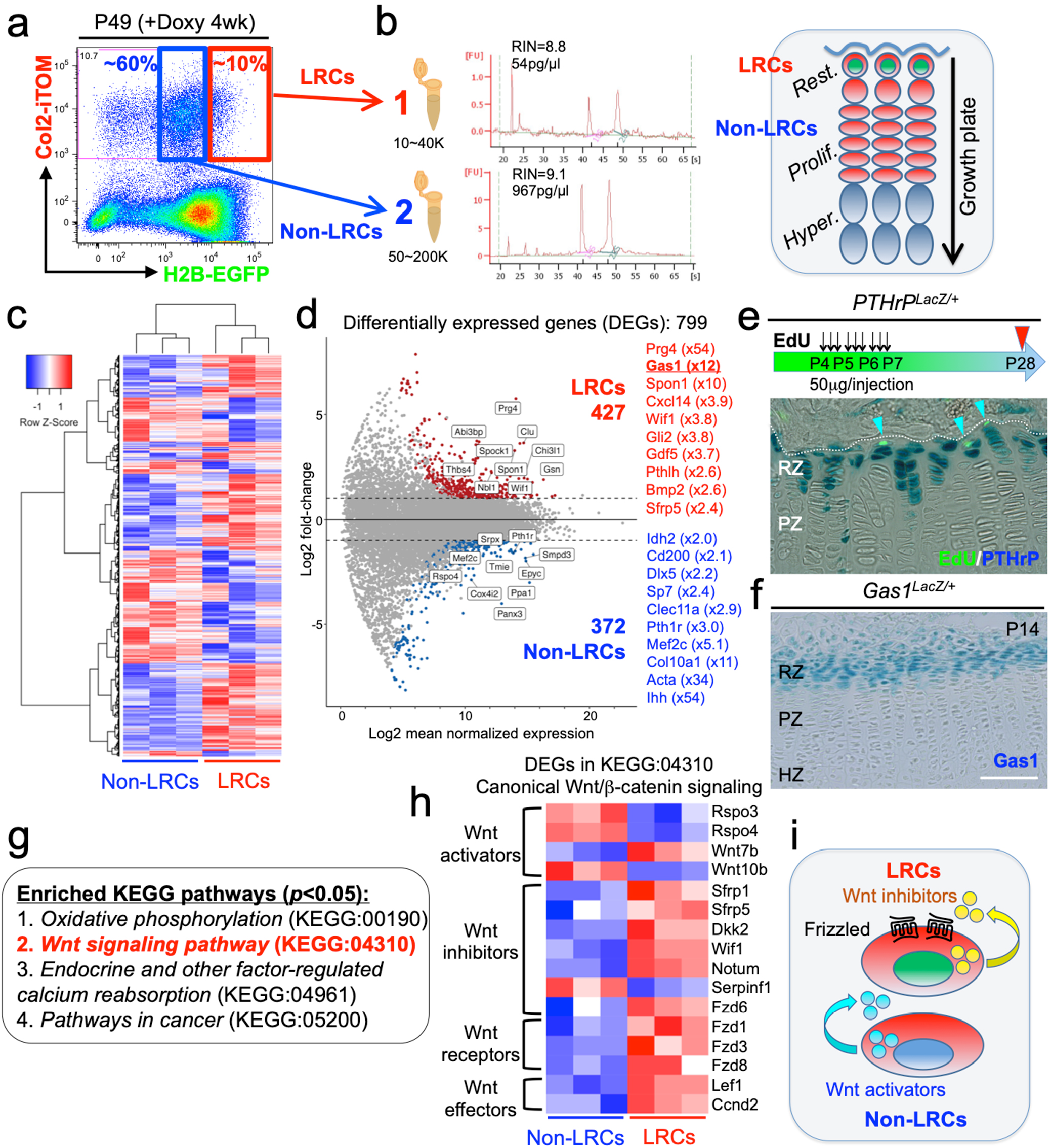
The unique molecular signature of label-retaining chondrocytes (LRCs) in the growth plate. **(a)** Cell sorting strategy to isolate LRCs (1: red box) and non-LRCs (2: blue box) after the chase, at P49 (+Doxy 4wk). **(b)** RNA integrity number (RIN) enumerated from bioanalyzer traces (28S/18S) of LRCs (top) and non-LRCs (bottom). Cartoon representation of GFP^+^/tdTomato^+^ LRCs populating resting zone and GFP^-^; tdTomato^+^ non-LRCs populating proliferating zone of growth plate (right). **(c)** Heatmap of top 500 differentially expressed genes (DEGs) with hierarchical clustering, between isolated non-LRCs and LRCs. *n*=3 biological replicates (i.e. three independent littermates of mice). **(d)** MA plot (Log2 fold change) of differentially expressed genes (DEGs) between isolated non-LRCs (372 total) and LRCs (427 total) with representative upregulated genes in each cell population. **(e)** *PTHrP*^*LacZ/+*^ distal femur growth plates with EdU administration, serially pulsed 9 times between P4 and P6 and analyzed after 22 days of chase at P28. Arrowheads: EdU label-retaining LacZ^+^ cells. RZ: resting zone, PZ: proliferating zone. *n*=6 mice. **(f)** *Gas1*^*LacZ/+*^ distal femur growth plates at P14. RZ: resting zone, PZ: proliferating zone, HZ: hypertrophic zone. Scale bar: 100μm. *n*=2 mice. **(g)** Enriched KEGG pathway terms (*p*<0.05) based on 799 differentially expressed genes (DEGs). **(h)** Heatmap of differentially expressed genes (DEGs) related to KEGG:04310 (canonical Wnt/β-Catenin signaling pathway). The DEGs were further classified by their functions in Wnt/β-Catenin signaling (e.g. Wnt activators, Wnt inhibitors, Wnt receptors and Wnt effectors). *n*=3 biological replicates (i.e. three independent littermates of mice). **(i)** Schematic diagram of Wnt activation and inhibition in non-LRCs and LRCs, respectively.

We further set out to validate the LRC markers using several independent approaches. First, to test if PTHrP^+^ cells overlap with LRCs, we performed an EdU pulse-chase experiment by serially pulsing *PTHrP-LacZ* knock-in mice (X. Chen et al., 2006) (Fig. S2a). Shortly after the pulse at P7, PTHrP^+^ cells were preferentially localized in an EdU-low zone, wherein 17.9±2.7% of EdU^+^ cells were PTHrP^+^ (Fig. S2a, left panel, and Fig. S2b). After 22 days of chase at P28, a great majority of EdU-retaining cells were PTHrP^+^, wherein 77.6±9.6% of EdU^+^ cells were PTHrP^+^ (Fig. 2e, Fig. S2a, right panel, and Fig. S2b). Therefore, LRCs become increasingly enriched among PTHrP^+^ chondrocytes in the postnatal growth plate. Second, we validated expression of a novel LRC marker, *Gas1*. Analysis of *Gas1-LacZ* knock-in mice (Martinelli & Fan, 2007) at P14 revealed that Gas1^+^ cells were exclusively found at the top of the growth plate corresponding to the resting zone (Fig. 2f). Therefore, *in vivo* expression patterns of two representative LRC markers – *PTHrP* and *Gas1* – using knock-in reporter lines further support the validity of the gene expression profile of LRCs that accurately reflects that of the resting zone of the growth plate.

Pathway analysis of DEGs revealed significant enrichment of four KEGG terms (*p*<0.05, FDR), including *Oxidative phosphorylation* (KEGG:00190), *Wnt signaling pathway* (KEGG:04310), *Endocrine and other factor-regulated calcium reabsorption* (KEGG:04961) and *Pathways in cancer* (KEGG:05200) (Fig. 2g). Notably, all DEGs annotated under the *Oxidative phosphorylation* KEGG term were upregulated in non-LRCs, highlighting a biochemically unique feature of non-LRCs undergoing active processes such as cell division and differentiation. Out of 21 DEGs annotated in *Wnt signaling pathway*, 16 genes were relevant to the canonical Wnt/β-catenin signaling pathway (Komiya & Habas, 2008) (Fig. 2h). Interestingly, LRCs were enriched for genes encoding canonical Wnt inhibitors such as *Sfrp1*, *Sfrp5*, *Dkk2*, *Wif1*, *Notum* and *Fzd6*, as well as genes encoding Wnt receptors such as *Fzd1*, *Fzd3* and *Fzd8*. Conversely, non-LRCs were enriched for genes encoding canonical Wnt activators such as *Rspo3*, *Rspo4*, *Wnt7b* and *Wnt10b* (Fig. 2h). Therefore, these RNA-seq analyses demonstrate that LRCs reside in a microenvironment in which inhibitors for canonical Wnt signaling are abundantly present in the milieu (Fig. 2i).

## 3. Activation of canonical Wnt signaling impairs formation, expansion and differentiation of PTHrP^+^ resting chondrocytes

We next set out to define how canonical Wnt signaling regulates slow-cycling chondrocytes of the postnatal growth plate. For this purpose, we activated Wnt/β-catenin signaling in PTHrP^+^ resting chondrocytes by conditionally inducing haploinsufficiency of *adenomatous polyposis coli (Apc)*, which is a critical component of the β-catenin degradation complex, using a *Pthrp-creER* (Mizuhashi et al., 2018) line and *Apc*-floxed allele (Cheung et al., 2010), and simultaneously traced the fates of these Wnt-activated PTHrP^+^ cells using an *R26R-*tdTomato reporter allele (Fig. 3a,b). Littermate triple transgenic mice with two corresponding genotypes – *PTHrP-creER*; *Apc*^+/+^; *R26R*^*tdTomato*^ (Control, PTHrP^CE^APC^++^ cells) and *PTHrP-creER*; *Apc*^fl/+^; *R26R*^*tdTomato*^ (APC cHet, PTHrP^CE^Apc^Het^ cells) mice – were pulsed with tamoxifen (250 μg) at P6 and analyzed at five consecutive time points after the chase, i.e. P9, P12, P21 P36 and P96 (Fig. 3c). Immunohistochemical analysis revealed that the β-catenin protein was upregulated in the resting zone of APC cHet growth plates and PTHrP^CE^tdTomato^+^ cells therein (Fig. 3d, leftmost panels, arrows), indicating that *Apc* haploinsufficiency indeed slowed β-catenin degradation and activated canonical Wnt signaling specifically in the resting zone of the growth plate.

**Figure 3.**
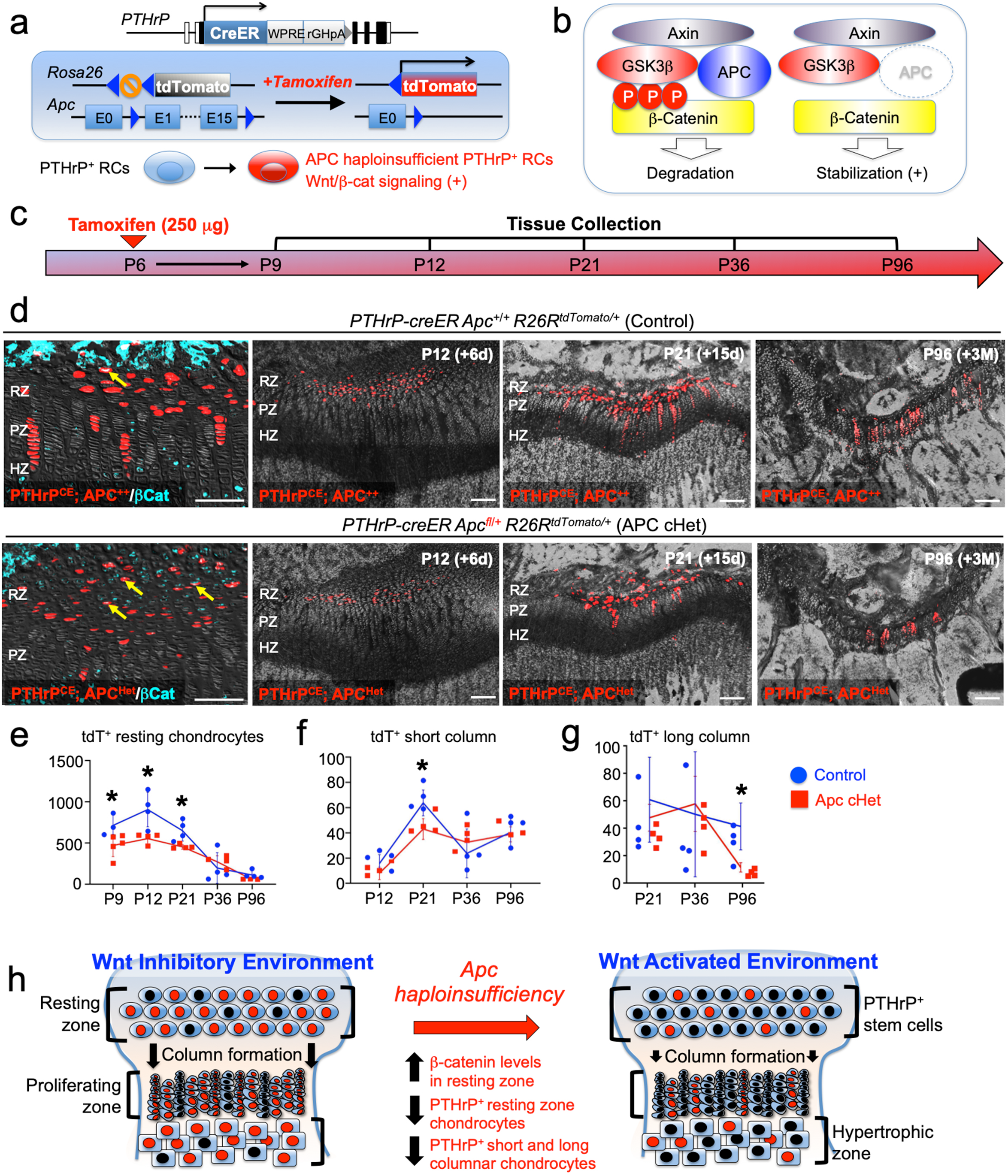
Activation of canonical Wnt/β-catenin signaling causes failure of formation and differentiation of PTHrP^+^ chondrocytes. **(a)** *PTHrP-creER*; *R26R*^*tdTomato*^ lineage-tracing model crossed with an *adenomatous polyposis coli (Apc)* floxed allele (flanking exons 1 and 15). Single intraperitoneal injection of tamoxifen (0.25 mg) at P6 induces *cre* recombination, leading to activation of canonical Wnt/β-catenin signaling in PTHrP^+^ chondrocytes via *Apc* haploinsufficiency (*PTHrP-creER; APC^fl/+^; R26R-tdTomato*). **(b)** Schematic diagram of β-catenin degradation complex. Phosphorylation of β-catenin protein leads to degradation (left). *Apc* haploinsufficiency leads to β-catenin stabilization by impairing the degradation complex (right). **(c)** Timeline for pulse-chase experiment. Tamoxifen injection (0.25 mg) at P6 and chase to P9, P12, P21, P36 and P96. **(d)** (Leftmost panel): β-catenin staining in *PTHrP-creER*; *Apc*^+/+^; *R26R*^*tdTomato*^ (Control) and *PTHrP-creER*; *Apc*^fl/+^; *R26R*^*tdTomato*^ (APC cHet) distal femur growth plates at P15. Arrows: β-catenin^+^tdTomato^+^ cells in RZ. (2^nd^-4^th^ panels): Distal femur growth plates of *PTHrP-creER*; *Apc*^+/+^; *R26R*^*tdTomato*^ (Control) and *PTHrP-creER*; *Apc*^fl/+^; *R26R*^*tdTomato*^ (APC cHet) at P12, P21 and P96. RZ: resting zone, PZ: proliferating zone, HZ: hypertrophic zone. Blue: β-catenin-Alexa633, red: tdTomato, gray: DAPI and DIC. Scale bars: 100 μm. *n*=4 mice per genotype per time point. **(e-g)** Compiled quantification data of total numbers of (e) resting chondrocytes, (f) short columnar chondrocytes (≤10 tdTomato^+^ cells) and (g) long columnar chondrocytes (>10 tdTomato^+^ cells) (P9: *n*=3 mice for Control, *n*=5 mice for Apc cHet, P12–P36: *n*=4 mice per genotype, P96: *n*=4 mice for Control, *n*=3 mice for Apc cHet), collected from serial sections of femur growth plates (2 femurs/mouse) at all time points. Asterisks represent significant differences between control and mutant groups based on *p*<0.05 using a Mann-Whitney’s *U*-test. Data are presented as mean ± s.d. Control versus Apc cHet, resting chondrocytes; P9: *p*=0.036, mean difference = 243.9±97.4, 95% confidence interval (4.2, 483.5); P12: *p*=0.029, mean difference = 351.9±109.8, 95% confidence interval (83.3, 620.5); P21: *p*=0.029, mean difference = 198.5±63.9, 95% confidence interval (42.1, 355.0); P36: *p*=0.343, mean difference = –76.3±100.3, 95% confidence interval (−321.8, 169.3); P96: *p*=0.057, mean difference = 55.3±28.7, 95% confidence interval (−18.5, 129.1). Control versus Apc cHet, short columns; P12: *p*=0.020, mean difference = 7.9±4.3, 95% confidence interval (−2.7, 18.5); P21: *p*=0.029, mean difference = 20.8±6.5, 95% confidence interval (5.0, 36.5); P36: *p*=0.343, mean difference = – 8.9±10.7, 95% confidence interval (−35.0, 17.3); P96: *p*=0.343, mean difference = 1.3±7.2, 95% confidence interval (−17.2, 19.7). Control versus Apc cHet, long columns; P21: *p*=0.886, mean difference = 10.0±12.1, 95% confidence interval (−19.6, 39.6); P36: *p*=0.686, mean difference = – 5.9±18.6, 95% confidence interval (−51.3, 39.5); P96: *p*=0.029, mean difference = 22.3±6.5, 95% confidence interval (6.2, 38.3). **(h)** PTHrP^+^ chondrocytes are maintained in a Wnt inhibitory environment within the resting zone. *Apc* haploinsufficiency increases β-catenin level in the resting zone, and decrease formation of PTHrP^+^ chondrocytes and their differentiation to columnar chondrocytes.

Subsequently, we quantified the numbers of lineage-marked tdTomato^+^ cells in the resting zone, as well as short (composed of ≤ tdTomato^+^ 10 cells) and long (composed of >10 tdTomato^+^ cells) columns of tdTomato^+^ chondrocytes based on serial sections of femur growth plates (Fig. 3d, right panels). Consistent with our prior report (Mizuhashi et al., 2018), PTHrP^CE^APC^++^ Control chondrocytes transiently increased in the resting zone during the first week of chase, and decreased thereafter due to the formation of columnar chondrocytes (P9: 718.7±132.7, P12: 910.3±209.9, P21: 655.4±125.0, P36: 200.3±187.2; P96: 116.1±48.5 cells, Fig. 3e, blue line, *n*=3–4 mice). In contrast, PTHrP^CE^APC^Het^ resting chondrocytes did not increase in number during the initial stage of chase, the numbers of which were significantly lower than those of Control at the initial three time points (P9: 474.8±134.8 [*p*=0.04], P12: 558.4±64.3 [*p*=0.03], P21: 443.4±79.2 [*p*=0.03] cells, Fig. 3e, red line, *n*=4–5 mice), and fell to levels that were similar to those in the Control at the latter two time points (Fig. 3d, rightmost panel, Fig. 3e). These data indicate that the formation and the expansion of PTHrP^+^ cells in the resting zone are impaired when canonical Wnt signaling is activated in these cells.

As expected, PTHrP^CE^APC^++^ resting chondrocytes established short columns (fewer than 10 cells/column) of tdTomato^+^ chondrocytes across the growth plate that peaked at P21 (P12: 20.0±7.1, P21: 67.4±10.1, P36: 27.5±19.4, P96=44.3±11.1 tdTomato^+^ columns, Fig. 3f, blue line, *n*=4 mice). The number of tdTomato^+^ short columns in APC cHet growth plates was reduced at P21 (P21: 45.9±7.7 tdTomato^+^ columns, Fig. 3f, red line, *n*=4 mice [*p*=0.03]), indicating that the formation of short columnar chondrocytes in the proliferating zone is inhibited upon canonical Wnt signaling activation. We suspect that this result reflects the reduction of PTHrP-creER^+^ cells in the resting zone in the preceding stages, though we cannot rule out direct effects of APC haploinsufficiency in the proliferating zone as well.

PTHrP^+^ resting chondrocytes continue to generate long columns (more than 10 cells/column) of tdTomato^+^ chondrocytes in the long term, the number of which gradually decreases until six months and reaches a plateau thereafter (Mizuhashi et al., 2018). Accordingly, PTHrP^CE^APC^++^ cells generated gradually decreasing but still substantial numbers of tdTomato^+^ long columns after 3 months of chase at P96 (P21: 44.4±23.2, P36: 36.1±34.1, P96: 26.5±12.4 tdTomato^+^ columns, Fig. 3g, blue line, *n*=4 mice). In contrast, the number of tdTomato^+^ long columns in APC cHet growth plates was significantly decreased at P96 (P96: 7.3±2.5 tdTomato^+^ columns, Fig. 3g, red line, *n*=4 mice [*p*=0.03]). Therefore, the ability for PTHrP^+^ resting chondrocytes to clonally establish columnar chondrocytes is impaired in response to activation of canonical Wnt signaling in the resting zone.

Taken together, these findings indicate that activation of canonical Wnt signaling impairs formation, expansion and differentiation of PTHrP^+^ chondrocytes in the resting zone (Fig. 3h). Thus, PTHrP^+^ resting chondrocytes are required to be maintained in a Wnt-inhibitory environment to maintain themselves and their column-forming capabilities.

## Discussion

In this study, we investigated the molecular mechanisms regulating the maintenance and the differentiation of slow-cycling chondrocytes localized in the resting zone of the postnatal growth plate. To date, our understanding of the molecular regulators of this special subclass of chondrocytes is largely grounded in histological and immunohistochemical observations and extrapolations from conditional gene ablation studies (Hallett et al., 2019). To address this gap in knowledge, we established a quadruple transgenic murine reporter model, “Col2-Q” system, to genetically label slow-cycling chondrocytes in an unbiased manner using a pulse-chase approach based on a chondrocyte-specific doxycycline-controllable Tet-Off system regulating expression of histone 2B-linked GFP. We successfully isolated label-retaining chondrocytes (LRCs) and their proliferating counterparts (non-LRCs) to profile the transcriptome of these cells. As the resting zone of the growth plate is considered to represent a resident stem-cell niche (Abad et al., 2002; Mizuhashi et al., 2018; Newton et al., 2019), our experiments serve as an approach to interrogate the fundamental characteristics of one of the stem-like cells residing in the postnatal growth plate.

It is unclear how slow-cycling chondrocytes in the resting zone maintain low mitotic capabilities while differentiating into columnar chondrocytes in the proliferating zone. Using a comparative bulk RNA-seq transcriptomic analysis, we discovered that LRCs are enriched for a unique set of genes associated with hallmark (e.g. *Pthrp* and *Sfrp5*) and novel (e.g. *Gas1*, *Spon1* and *Wif1*) markers for resting chondrocytes, in addition to Wnt inhibitory molecules (e.g. *Sfrp1*, *Dkk2*, *Notum* and *Fzd6*). Conversely, non-LRCs were enriched for markers of pre-hypertrophic (e.g. *Ihh*) and hypertrophic (e.g. *Col10a1*) chondrocytes, and represent differentially expressed genes commonly associated with metabolically active cellular processes, such as oxidative phosphorylation. We further validated the expression of *Pthlh*, which is a hallmark marker for resting chondrocytes, and *Gas1*, a novel marker, using *PTHrP-LacZ* and *Gas1-LacZ* knock-in reporter alleles, respectively. We found that PTHrP^*+*^ chondrocytes in the resting zone maintain low levels of mitotic activity, indicated by EdU labeling and pulse-chase experiments. Thus, the genes identified by our comparative transcriptomic analysis appear to represent accurate transcriptomic features of distinct populations of slow-cycling versus differentiating chondrocytes in the postnatal growth plate. Future investigations aimed at assessing the roles of novel marker genes may lead to the identification of novel skeletal stem cell populations that are important for the postnatal growth plate.

Wnt/β-catenin signaling is essential for endochondral ossification (Regard et al., 2012), and is shown to regulate initiation of chondrocyte hypertrophy by inhibiting PTHrP signaling activities (Guo et al., 2009). Moreover, Wnt/β-catenin signaling is essential during skeletal development for regulating mesenchymal progenitor differentiation in favor of osteoblasts (Day et al., 2005), or for preventing transdifferentiation of osteoblast precursors into chondrocytes (Hill et al., 2005). However, no previous report ties Wnt signaling to the maintenance of putative skeletal stem cell populations in the resting zone of the growth plate. In order to determine the functional contribution of Wnt signaling to PTHrP^+^ resting chondrocyte skeletal stem cells and their differentiation, one copy of *adenomatous polyposis coli* (*Apc*), a critical signaling component of the β-catenin degradation complex, was selectively ablated using a resting chondrocyte-specific *Pthrp-creER* line. In the resting zone, Apc haploinsufficiency led to increased β-catenin protein expression specifically in the resting zone including in PTHrP^+^ chondrocytes, decreased formation and expansion of PTHrP^+^ chondrocytes, and reduced differentiation capabilities of these cells into columnar chondrocytes in the proliferating zone. Therefore, canonical Wnt signaling plays an important role in modulating PTHrP^+^ chondrocytes in the resting zone and regulating their differentiation.

Taken together, our data support a novel paradigm that slow-cycling PTHrP^+^ chondrocytes are maintained in a canonical Wnt-inhibitory environment within the resting zone of the growth plate, and that this relationship is critical to regulating the formation, the expansion and the differentiation of chondrocytes of the resting zone.

## Materials and Methods

### Generation of *Col2a1-tTA* transgenic mice

*Col2a1-tTA* transgenic mice were generated by pronuclear injection of a NotI-digested 8.4kb gene construct containing a 3kb mouse *Col2a1* promoter and a 3kb fragment of intron 1 ligated to a splice acceptor sequence followed by an internal ribosome-entry site (IRES) (Ovchinnikov et al., 2000), tetracycline-controlled transactivator (tTA) and the SV40 large T antigen polyadenylation signal (Takara Bio, Mountain View, CA), into B6SJLF1 fertilized eggs. The G0 founder mice were backcrossed with C57/BL6 mice at least for three generations. Of the two lines established, the high expresser line (Line H) was used for subsequent studies. The insertion site of the *Col2a1-tTA* transgene was determined based on the Genome Walker Universal system (Takara Bio). The *Col2a1-tTA* transgene was inserted 16kbp downstream of *Pellino2* on Chromosome 14. *Col2a1-tTA* mice were genotyped using PCR primers discriminating heterozygosity and homozygosity of the transgene (85: SV40pA_End_Fw: ACGGGAAGTATCAGCTCGAC, 86: Mm14_5WT_Fw: TTGAGAGTCTCCCAGGCAAT, 87: Mm14_3WT_Rv: CTCCTGATCTCCTGGCAAAG, ~600bp for wild-type, ~300bp for Col2a1-tTA allele).

### Mice

*TRE-H2B-EGFP* (Foudi et al., 2009) knock-in, *Col2a1-creER* transgenic (Nakamura et al., 2006), *PTHrP-LacZ/null* knock-in (X. Chen et al., 2006), *Gas1-LacZ/null* knock-in (Martinelli & Fan, 2007), *PTHrP-creER* transgenic (Mizuhashi et al., 2018) mice have been described elsewhere. *Rosa26-CAG-loxP-stop-loxP-*tdTomato (Ai14: *R26R*-tdTomato, JAX007914), *Apc*-floxed (JAX009045) mice (Cheung et al., 2010) were acquired from the Jackson Laboratory. All procedures were conducted in compliance with the Guidelines for the Care and Use of Laboratory Animals approved by the University Michigan’s Institutional Animal Care and Use Committee (IACUC), protocol 7681 and 9496. All mice were housed in a specific pathogen-free condition, and analyzed in a mixed background. Mice were identified by micro-tattooing or ear tags. Tail biopsies of mice were lysed by a HotShot protocol (incubating the tail sample at 95°C for 30 min in an alkaline lysis reagent followed by neutralization) and used for GoTaq Green Master Mix PCR-based genotyping (Promega, and Nexus X2, Madison, WI). Mice were euthanized by over-dosage of carbon dioxide or decapitation under inhalation anesthesia in a drop jar (Fluriso, Isoflurane USP, VetOne, Boise, ID).

### Doxycycline

Mice were weaned at postnatal day (P) 21 and fed with a standard diet containing 2mg/g doxycycline (Bio-Serv F3893, Flemington, NJ) for up to 9 weeks.

### Tamoxifen

Tamoxifen (Sigma T5648, St. Louis, MO) was mixed with 100% ethanol until completely dissolved. Subsequently, a proper volume of sunflower seed oil (Sigma S5007) was added to the tamoxifen-ethanol mixture and rigorously mixed. The tamoxifen-ethanol-oil mixture was incubated at 60°C in a chemical hood until the ethanol evaporated completely. The tamoxifen-oil mixture was stored at room temperature until use. Mice with 21 days of age or older received two doses of 2mg of tamoxifen intraperitoneally at 3 and 2 days prior to analysis, or mice with 6 days of age received a single dose of 0.25mg tamoxifen intraperitoneally for lineage-tracing analysis.

### Cell proliferation and EdU label-retention assay

5-ethynyl-2’-deoxyuridine (EdU) (Invitrogen A10044, Carlsbad, CA) dissolved in PBS was administered to mice at indicated postnatal days. Click-iT Imaging Kit with Alexa Flour 488-azide (Invitrogen C10337) was used to detect EdU in cryosections. For EdU label-retention assay, *PTHrP-LacZ* mice received serial doses of EdU (50μg each) between P4 and P6, and chased for 3 weeks.

### X-Gal staining of dissected femur epiphyses

Distal epiphyses of femurs were manually dislodged, and attached soft tissues were carefully removed to ensure the maximum penetration of the substrate. Dissected epiphyses were fixed in 2% paraformaldehyde for 30 min. at 4C°, followed by overnight X-gal staining at 37°C. Stained samples were further postfixed in 4% paraformaldehyde, overnight at 4°C, then decalcified in 15% EDTA for 7 days. Decalcified samples were cryoprotected in 30% sucrose/PBS followed by 30% sucrose/PBS:OCT (1:1) solution, each overnight at 4°C.

### Histology

Bilateral femurs were dissected under a stereomicroscope (Nikon SMZ-800, Minato City, Japan) to remove soft tissues, and fixed in 4% paraformaldehyde for a proper period, typically ranging from 3 hours to overnight at 4°C, then decalcified in 15% EDTA for a proper period, typically ranging from 0 hours to 14 days. Decalcified samples were cryoprotected in 30% sucrose/PBS solutions and then in 30% sucrose/PBS:OCT (1:1) solutions, each at least overnight at 4°C. Samples were embedded in an OCT compound (Tissue-Tek, Sakura, Torrance, CA) under a stereomicroscope and transferred on a sheet of dry ice to solidify the compound. Embedded samples were cryosectioned at 14–50μm using a cryostat (Leica CM1850, Wetzlar, Germany) and adhered to positively charged glass slides (Fisherbrand ColorFrost Plus). Cryosections were stored at –20°C (quantification) or –80°C (immunofluorescence) in freezers until use. Sections were postfixed in 4% paraformaldehyde for 15 min at room temperature. For functional conditional knockout experiments, 50μm serial sections were collected through the entire growth plate. For immunofluorescence experiments, epiphyses were popped out of bilateral femurs, processed for 24 hours in 4% paraformaldehyde and sectioned at 14μm. Sections were incubated with anti-β-catenin primary antibody (Abcam ab16051, Cambridge, UK) overnight at 4°C and further stained with 1:200 Alexa Fluor 633 Goat anti-Rabbit IgG (H+L) Secondary Antibody (Invitrogen A21071) at a 20°C for 3 hours. Sections were further incubated with DAPI (4’,6-diamidino-2-phenylindole, 5μg/ml, Invitrogen D1306) to stain nuclei prior to imaging. For EdU assay, sections were incubated with Alexa Fluor 488-azide (Invitrogen A10266) for 30 min at 43°C using Click-iT Imaging Kit (Invitrogen C10337). Sections were further incubated with DAPI to stain nuclei prior to imaging. Stained samples were mounted in TBS with No.1.5 coverslips (Fisher, Waltham, MA).

### Imaging and cell quantification

Images were captured by a fluorescence microscope (Nikon Eclipse E800) with prefigured triple-band filter settings for DAPI/FITC/TRITC, and merged with Spot Advanced Software (Spot Imaging, Sterling Heights, MI), or an automated inverted fluorescence microscope with a structured illumination system (Zeiss Axio Observer Z1 with ApoTome.2 system) and Zen 2 (blue edition) software. Confocal images were acquired using LSM510 and Zen2009 software (Zeiss, Oberkochen, Germany) with lasers and corresponding band-pass filters for DAPI (Ex.405nm, BP420-480), GFP (Ex.488nm, BP505-530) and tdTomato (Ex.543nm, BP565-595).

LSM Image Viewer and Adobe Photoshop software were used to capture and align images. Cells were counted by two individuals using single blinded methods to ensure unbiased data interpretation.

### Growth plate cell preparation

Distal epiphyses of femurs and proximal epiphyses of tibias were manually dislodged using dull scissors, and attached soft tissues and woven bones were carefully removed using a cuticle nipper. Cells were dissociated from dissected epiphyses using five serial rounds of collagenase digestion, incubating with 2 Wunsch units of Liberase TM (Roche, Basel, Switzerland) in 2ml Ca^2+^, Mg^2+^-free Hank’s Balanced Salt Solution (HBSS, Sigma H6648) at 37°C for 30 min. each time on a shaking incubator (ThermomixerR, Eppendorf, Hamburg, Germany). Single cell suspension was generated using an 18-gauge needle and a 1ml Luer-Lok syringe (BD), and filtered through a 70μm cell strainer (BD) into a 50ml tube on ice.

### Flow cytometry

Dissociated cells were stained by standard protocols with allophycocyanin (APC)-conjugated anti-mouse CD45 (30F-11) antibodies (1:500, eBioscience, San Diego, CA). Flow cytometry analysis was performed using a four-laser BD LSR II (Ex. 355/407/488/633 nm) and FACSDiva software. Acquired raw data were further analyzed on FlowJo software (TreeStar). Representative plots of at least three independent biological samples are shown in the figures.

### Fluorescence-activated cell sorting (FACS) and RNA isolation

Cell sorting was performed using a five-laser BD FACS Aria II (Ex.355/407/488/532/633nm) and FACSDiva. CD45^neg^GFP^high^ cells at the top 10 percentile of GFP brightness (LRCs) and CD45^neg^GFP^mid-low^ cells with 10~70 percentile of GFP brightness (non-LRCs) were directly sorted into TRIzol LS Reagent (ThermoFisher 10296010, Waltham, MA). Total RNA was isolated using NucleoSpin RNA XS (Macherey-Nagel, 740902). RNA Integrity Number (RIN) was assessed by Agilent 2100 Bioanalyzer RNA 6000 Pico Kit. Samples with RIN>8.0 were used for subsequent analyses.

### RNA amplification and deep sequencing

Complementary DNAs were prepared by SMART-Seq v4 Ultra Low Input RNA Kit for Sequencing (Takara 634888) using 150~800pg of total RNA. Post-amplification quality control was performed by Agilent TapeStation DNA High Sensitivity D1000 Screen Tape system. DNA libraries were prepared by Nextera XT DNA Library Preparation Kit (Illumina) and submitted for deep sequencing (Illumina HiSeq 2500).

### RNA-seq analysis

cDNA libraries were sequenced using following conditions; six samples per lane, 50 cycle single end read. Reads files were downloaded and concatenated into a single .fastq file for each sample. The quality of the raw reads data for each sample was checked using FastQC to identify quality problems. Tuxedo Suite software package was subsequently used for alignment (using TopHat and Bowtie2), differential expression analysis, and post-analysis diagnostics. FastQC was used for a second round of quality control (post-alignment). HTSeq/DESeq2 was run using UCSC mm10.fa as the reference genome sequence. Expression quantitation was performed with HTSeq, to count non-ambiguously mapped reads only. HTSeq counts per gene were then used in a custom DESeq2 paired analysis. Normalization and differential expression were performed with DESeq2, using a negative binomial generalized linear model, including a term for mouse of origin for a paired analysis. Plots were generated using variations or alternative representations of native DESeq2 plotting functions, ggplot2, plotly, and other packages within the R environment. Heatmaps were generated with updated rlog normalized count values for each sample for all plus top sets (500) of differentially expressed genes with the gplots package (v 3.0.1). Two types of clustering were used: 1) averaging across rows with Pearson correlation distance with average linkage and 2) Ward’s squared dissimilarity criterion. Top differentially expressed genes were determined after ranking genes by standard deviation across all samples. Independent of iPathway, GO term enrichment was performed on DE results, with a logFC threshold of 2 and adjusted *p*-value < 0.05 with the GOseq package (v 1.36) with probability weighting function and GO enrichment specified with mm10 as genome and gene symbol specified as gene ID format. Results were plotted for the top ten of selected terms related to the Wnt pathways, ranked by overrepresented p-value using ggplot2 (v 3.2.1). KEGG results with FDR correction and gene tables for Wnt signaling pathway were downloaded from iPathway (report ID: 41865). KEGG gene tables for each pathway were used to subset the DE results before restricting results to genes for which both log fold change and adjusted *p*-value statistics were available.

### Replicates

All experiments were performed in biological replicates. For all data presented in the manuscript, we examined at least three independent biological samples (three different mice) to ensure the reproducibility. Biological replicates were defined as multiple experimental samples sharing common genotypes and genetic backgrounds. For each series of the experiments, all attempts at biological replication were successful. Technical replicates were generated from a single experimental sample. For example, serial sections of the femur growth plate from a single mouse were considered technical replicates. Outliers were uncommon in our datasets and did not impact the trend and the significance of our quantitated results. As a result, all quantitative data were included to ensure transparency in our data interpretation.

### Statistical analysis

Results are presented as mean values ± S.D. Statistical evaluation was conducted based on Mann-Whitney’s *U*-test. A *p* value <10.05 was considered significant. No statistical method was used to predetermine sample size. Sample size was determined on the basis of previous literature and our previous experience to give sufficient standard deviations of the mean so as not to miss a biologically important difference between groups. The experiments were not randomized. All of the available mice of the desired genotypes were used for experiments. The investigators were not blinded during experiments and outcome assessment. One femur from each mouse was arbitrarily chosen for histological analysis. Genotypes were not particularly highlighted during quantification.

## Data availability

The bulk RNA-seq datasets presented herein have been deposited in the National Center for Biotechnology Information (NCBI)’s Gene Expression Omnibus (GEO), and are accessible through GEO Series accession numbers GSE160364 [https://www.ncbi.nlm.nih.gov/geo/query/acc.cgi?acc=GSE160364]. The source data underlying all Figures and Supplementary Figures are provided as a Source Data file. All the raw images and flow cytometry files supporting the conclusion of this study will be deposited in Dryad Digital Repository during the revision.

## Acknowledgements

This research was supported by grants from National Institute of Health (R01DE026666 to N.O., R03DE027421 to W.O., P01DK011794 to H.M.K. and T32DE007057 to S.A.H.).

We thank H. Hock for *TRE-H2B-EGFP* mice and B. Allen for *Gas1-LacZ* mice.

We acknowledge support from the Bioinformatics Core of the University of Michigan Medical School’s Biomedical Research Core Facilities.

---

**Figure S1.**
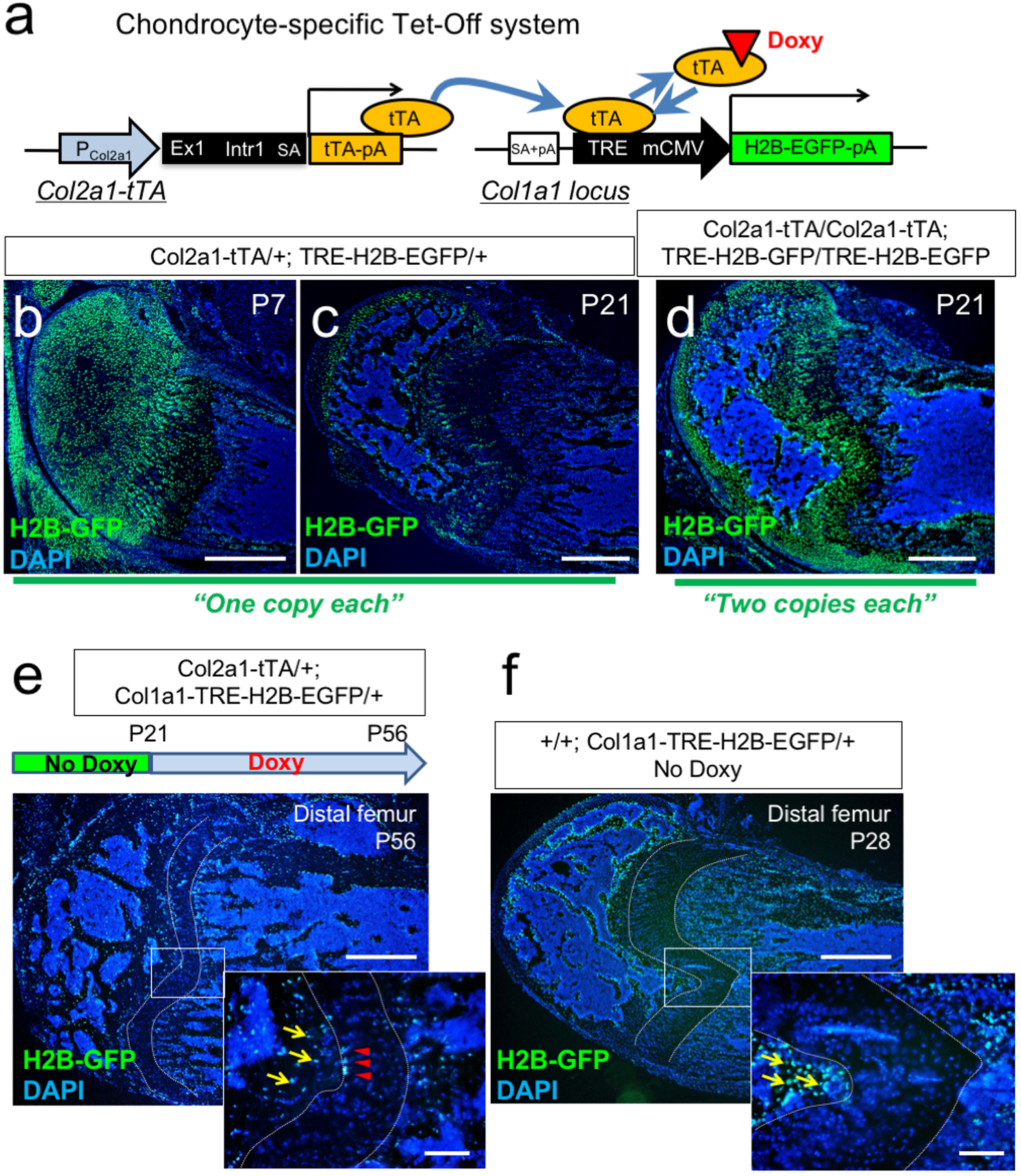
A genetic label-retention strategy to identify slow-cycling chondrocytes. **(a)** Chondrocyte-specific Tet-Off system by *Col2a1-tTA* and *TRE-H2B-EGFP* transgenes. During development, *Col2a1*^+^ cells accumulate H2B-EGFP in the nucleus. Binding of tetracycline-controlled transactivator (tTA) to Tet-responsive element (TRE) is prevented in the presence of doxycycline. As a result of this chase, slow­ cycling cells retain a high level of H2B-EGFP, while proliferating cells dilute H2B-EGFP signal as they continue to divide. **(b,c)** Distal femur growth plates of Col2a1-tTA/+; TRE-H2B-EGFP/+ double heterozygous mice at P? (b) and P21 (c). Note that only a small fraction of growth plates marked by GFP in (c). Scale bars: S00μm. *n=3* mice. **(d)** Distal femur growth plates of Col2a1-tTA/Col2a1-tTA; TRE-H2B-EGFP/TRE-H2B-EGFP double homozygous mice at P21. Note that a greater number of growth plate cells are marked by GFP than in (c). Scale bars: S00μm. *n=3* mice. **(e)** Distal femur growth plates of Col2a1-tTA/+; Col1a1-TRE-H2B-EGFP/+ mice, after 5 weeks of chase at P56. Arrowheads: Gf Phigh label-retaining chondrocytes, arrows: GfP+ osteoblasts/cytes. Scale bars: 500μm, 200μm (inset). *n= 3* mice. **(f)** Distal femur growth plates of +/+; Col1a1-TRE-H2B-EGFP/+ mice at P28. Arrows: Gf P+ osteoblasts/cytes. Scale bars: 500μm, 200μm (inset). *n= 3* mice.

**Figure S2.**
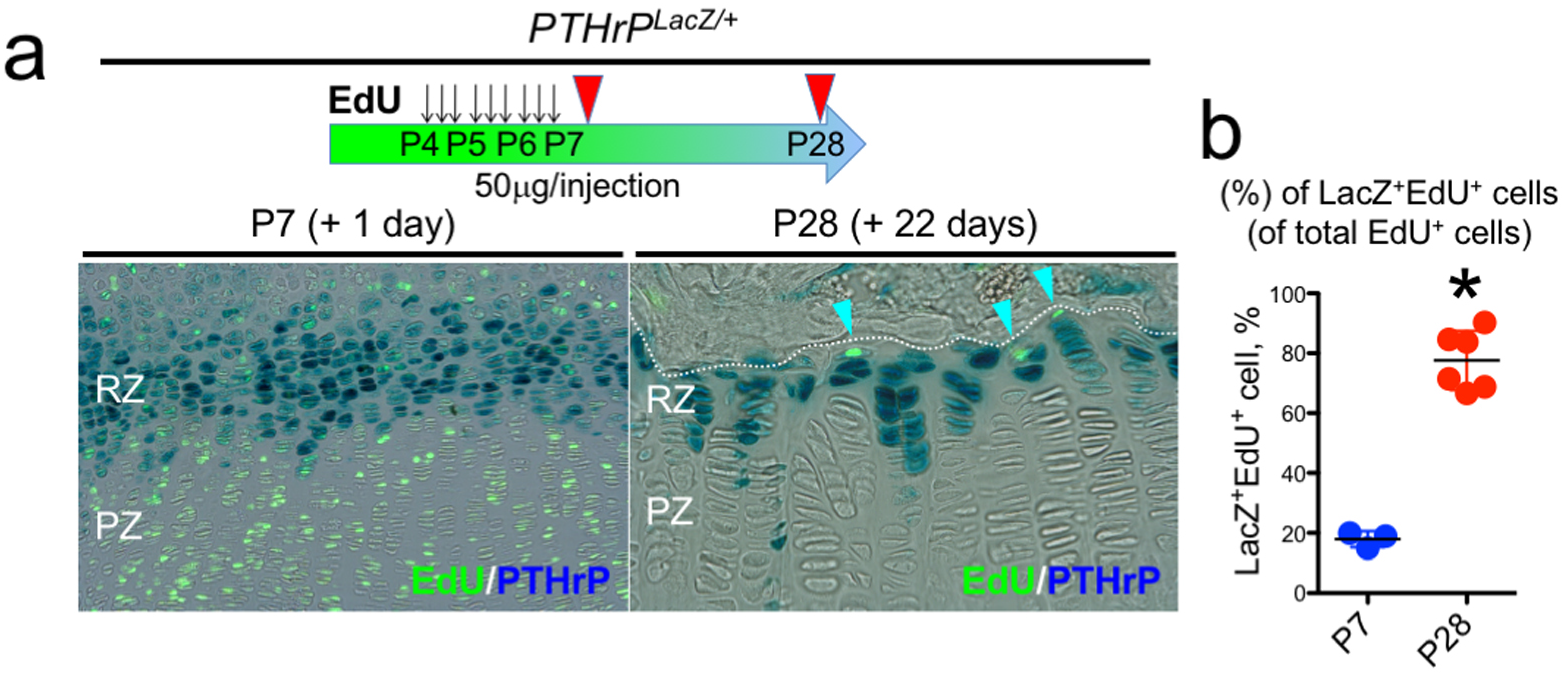
Label-retaining chondrocytes (LRCs) are enriched among PTHrP^+^ chondrocytes. **(a,b)** *PTHrP*^*LacZ/+*^ distal femur growth plates with EdU administration, serially pulsed 9 times between P4 and P6. (a, left panel): Immediately after the pulse at P7. (a, right panel): After 22 days of chase at P28. Arrowheads: EdU label-retaining LacZ^+^ cells. RZ: resting zone, PZ: proliferating zone. Scale bars: 100μm. (b): The percentage of LacZ^+^EdU^+^ cells among total EdU^+^ cells, at P7 *(n=3* mice) and P28 *(n=6* mice). *\*p<0.05,* Mann-Whitney’s *U-test.* Data are presented as mean ± s.d. P7 versus P28: p=0.024, mean difference = −59.7±6.0, 95% confidence interval (−73.8, −45.5).

